## Supplementary Materials for "Overcoming barriers to the registration of new varieties"

Chin Jian Yang

Joanne Russell

Luke Ramsay

William Thomas

Wayne Powell

Ian Mackay

**Table S1. DUS trait scoring system.** Currently, the UK barley DUS system includes 28 morphological traits that are scored based on discrete scales. The scales are different from each trait, but are generally ranging from 1 to 9 with few exceptions.

| Index | Trait | Score | Description |
|---|---|---|---|
| 1 | kernel: colour of aleurone layer | 1 | whitish |
| | | 2 | weakly coloured |
| | | 3 | strongly coloured |
| 2 | plant: growth habit | 1 | erect |
| | | 2 | erect to semi-erect |
| | | 3 | semi-erect |
| | | 4 | semi-erect to intermediate |
| | | 5 | intermediate |
| | | 6 | intermediate to semi-prostrate |
| | | 7 | semi-prostrate |
| | | 8 | semi-prostrate to prostrate |
| | | 9 | prostrate |
| 3 | lowest leaves: hairiness of leaf sheaths | 1 | absent |
| | | 9 | present |
| 4 | flag leaf: intensity of anthocyanin colouration of auricles | 1 | absent |
| | | 2 | absent to weak |
| | | 3 | weak |
| | | 4 | weak to medium |
| | | 5 | medium |
| | | 6 | medium to strong |
| | | 7 | strong |
| | | 8 | strong to very strong |
| | | 9 | very strong |
| 5 | flag leaf: attitude | 1 | erect |
| | | 2 | erect to semi-erect |
| | | 3 | semi-erect |
| | | 4 | semi-erect to horizontal |
| | | 5 | horizontal |
| | | 6 | horizontal to semi-drooping |
| | | 7 | semi-drooping |
| | | 8 | semi-drooping to drooping |
| | | 9 | drooping |
| 6 | flag leaf: glaucosity of sheath | 1 | absent |
| | | 2 | absent to weak |
| | | 3 | weak |
| | | 4 | weak to medium |
| | | 5 | medium |
| | | 6 | medium to strong |
| | | 7 | strong |
| | | 8 | strong to very strong |
| | | 9 | very strong |

| | | | |
|---|---|---|---|
| 7 | time of ear emergence (first spikelet visible on 50% of ears) | 1 | very early |
| | | 2 | very early to early |
| | | 3 | early |
| | | 4 | early to medium |
| | | 5 | medium |
| | | 6 | medium to late |
| | | 7 | late |
| | | 8 | late to very late |
| | | 9 | very late |
| 8 | awns: intensity of anthocyanin colouration of tips | 1 | absent |
| | | 2 | absent to weak |
| | | 3 | weak |
| | | 4 | weak to medium |
| | | 5 | medium |
| | | 6 | medium to strong |
| | | 7 | strong |
| | | 8 | strong to very strong |
| | | 9 | very strong |
| 9 | ear: glaucosity | 1 | absent |
| | | 2 | absent to weak |
| | | 3 | weak |
| | | 4 | weak to medium |
| | | 5 | medium |
| | | 6 | medium to strong |
| | | 7 | strong |
| | | 8 | strong to very strong |
| | | 9 | very strong |
| 10 | ear: attitude | 1 | erect |
| | | 2 | erect to semi-erect |
| | | 3 | semi-erect |
| | | 4 | semi-erect to horizontal |
| | | 5 | horizontal |
| | | 6 | horizontal to semi-recurved |
| | | 7 | semi-recurved |
| | | 8 | semi-recurved to recurved |
| | | 9 | recurved |
| 11 | plant: length (stem, ear and awns) | 1 | very short |
| | | 2 | very short to short |
| | | 3 | short |
| | | 4 | short to medium |
| | | 5 | medium |
| | | 6 | medium to long |
| | | 7 | long |
| | | 8 | long to very long |
| | | 9 | very long |
| 12 | ear: number of rows | 1 | two |
| | | 2 | more than two |

| | | | |
|---|---|---|---|
| 13 | ear: shape | 3 | tapering |
| | | 4 | tapering to parallel |
| | | 5 | parallel |
| | | 6 | parallel to fusiform |
| | | 7 | fusiform |
| 14 | ear: density | 1 | very lax |
| | | 2 | very lax to lax |
| | | 3 | lax |
| | | 4 | lax to medium |
| | | 5 | medium |
| | | 6 | medium to dense |
| | | 7 | dense |
| | | 8 | dense to very dense |
| | | 9 | very dense |
| 15 | ear: length (excluding awns) | 1 | very short |
| | | 2 | very short to short |
| | | 3 | short |
| | | 4 | short to medium |
| | | 5 | medium |
| | | 6 | medium to long |
| | | 7 | long |
| | | 8 | long to very long |
| | | 9 | very long |
| 16 | awn: length (compared to ear) | 3 | short |
| | | 4 | short to medium |
| | | 5 | medium |
| | | 6 | medium to long |
| | | 7 | long |
| 17 | rachis: length of first segment | 3 | short |
| | | 4 | short to medium |
| | | 5 | medium |
| | | 6 | medium to long |
| | | 7 | long |
| 18 | rachis: curvature of first segment | 1 | absent |
| | | 2 | absent to weak |
| | | 3 | weak |
| | | 4 | weak to medium |
| | | 5 | medium |
| | | 6 | medium to strong |
| | | 7 | strong |
| | | 8 | strong to very strong |
| | | 9 | very strong |
| 19 | ear: development of sterile spikelets | 1 | deficiens |
| | | 2 | full |
| 20 | sterile spikelets: attitude (in mid-third of ear) | 1 | parallel |
| | | 2 | parallel to weakly divergent |
| | | 3 | divergent |

| | | | |
|---|---|---|---|
| 21 | median spikelet: length of glume and its awn relative to grain | 1 | shorter |
| | | 2 | equal |
| | | 3 | longer |
| 22 | grain: rachilla hair type | 1 | short |
| | | 2 | long |
| 23 | grain: husk | 1 | absent |
| | | 9 | present |
| 24 | grain: anthocyanin colouration of nerves of lemma | 1 | absent |
| | | 2 | absent to weak |
| | | 3 | weak |
| | | 4 | weak to medium |
| | | 5 | medium |
| | | 6 | medium to strong |
| | | 7 | strong |
| | | 8 | strong to very strong |
| | | 9 | very strong |
| 25 | grain: spiculation of inner lateral nerves of dorsal side of lemma | 1 | absent |
| | | 2 | absent to weak |
| | | 3 | weak |
| | | 4 | weak to medium |
| | | 5 | medium |
| | | 6 | medium to strong |
| | | 7 | strong |
| | | 8 | strong to very strong |
| | | 9 | very strong |
| 26 | grain: hairiness of ventral furrow | 1 | absent |
| | | 9 | present |
| 27 | grain: disposition of lodicules | 1 | frontal |
| | | 2 | clasping |
| 28 | seasonal type | 1 | winter |
| | | 2 | alternative |
| | | 3 | spring |

**Table S2. Genetic variances, phenotypic variances and heritabilities for 28 DUS traits**. Estimates of genetic variance ($V_G$), phenotypic variance ($V_P$) and heritabilities ($h^2$) are shown along with their standard errors in parentheses.

| Trait | $V_G$ | | | $V_P$ | | | $h^2$ | | |
|---|---|---|---|---|---|---|---|---|---|
| | Combined | Spring | Winter | Combined | Spring | Winter | Combined | Spring | Winter |
| 1 | 0.22 (0.02) | 0.01 (0.00) | 0.31 (0.05) | 0.28 (0.02) | 0.07 (0.01) | 0.40 (0.04) | 0.78 (0.04) | 0.16 (0.06) | 0.79 (0.06) |
| 2 | 0.29 (0.06) | 0.19 (0.06) | 0.25 (0.08) | 1.16 (0.06) | 1.09 (0.08) | 1.04 (0.08) | 0.25 (0.05) | 0.17 (0.06) | 0.24 (0.07) |
| 3 | 1.86 (0.20) | NA | 2.96 (0.47) | 2.48 (0.16) | NA | 4.29 (0.37) | 0.75 (0.04) | NA | 0.69 (0.07) |
| 4 | 1.94 (0.23) | 0.19 (0.06) | 3.01 (0.49) | 2.62 (0.18) | 1.04 (0.08) | 3.59 (0.37) | 0.74 (0.05) | 0.19 (0.06) | 0.84 (0.08) |
| 5 | 0.91 (0.40) | 0.79 (0.49) | 0.68 (0.41) | 3.21 (0.37) | 2.83 (0.53) | 2.69 (0.44) | 0.28 (0.13) | 0.28 (0.19) | 0.25 (0.16) |
| 6 | 0.12 (0.04) | 0.03 (0.03) | 0.14 (0.07) | 1.04 (0.06) | 0.75 (0.06) | 1.34 (0.11) | 0.12 (0.04) | 0.05 (0.03) | 0.10 (0.05) |
| 7 | 0.53 (0.11) | 0.28 (0.09) | 0.56 (0.16) | 1.88 (0.10) | 1.41 (0.11) | 2.16 (0.17) | 0.28 (0.05) | 0.20 (0.06) | 0.26 (0.07) |
| 8 | 1.46 (0.18) | 0.10 (0.05) | 2.29 (0.38) | 2.17 (0.15) | 1.02 (0.08) | 2.78 (0.29) | 0.67 (0.05) | 0.09 (0.04) | 0.83 (0.08) |
| 9 | 0.65 (0.10) | 0.52 (0.10) | 0.54 (0.14) | 1.55 (0.09) | 1.14 (0.09) | 1.66 (0.13) | 0.42 (0.05) | 0.45 (0.07) | 0.33 (0.08) |
| 10 | 0.47 (0.10) | 0.36 (0.10) | 0.35 (0.13) | 1.86 (0.10) | 1.39 (0.10) | 2.12 (0.17) | 0.25 (0.05) | 0.26 (0.07) | 0.17 (0.06) |
| 11 | 0.27 (0.07) | 0.18 (0.07) | 0.23 (0.10) | 1.58 (0.09) | 1.42 (0.11) | 1.65 (0.13) | 0.17 (0.04) | 0.13 (0.05) | 0.14 (0.06) |
| 12 | 0.01 (0.00) | NA | 0.01 (0.00) | 0.01 (0.00) | NA | 0.01 (0.00) | 1.00 (0.01) | NA | 1.00 (0.03) |
| 13 | 0.08 (0.03) | 0.03 (0.03) | 0.07 (0.04) | 0.85 (0.05) | 0.86 (0.07) | 0.81 (0.07) | 0.10 (0.04) | 0.04 (0.03) | 0.09 (0.05) |
| 14 | 0.28 (0.06) | 0.15 (0.06) | 0.29 (0.09) | 1.23 (0.07) | 1.12 (0.08) | 1.24 (0.10) | 0.23 (0.05) | 0.14 (0.05) | 0.24 (0.07) |
| 15 | 0.12 (0.03) | 0.04 (0.03) | 0.17 (0.05) | 0.69 (0.04) | 0.74 (0.06) | 0.58 (0.05) | 0.18 (0.05) | 0.05 (0.04) | 0.29 (0.08) |
| 16 | 0.14 (0.04) | 0.08 (0.03) | 0.11 (0.05) | 0.80 (0.04) | 0.54 (0.04) | 1.01 (0.08) | 0.18 (0.04) | 0.15 (0.05) | 0.11 (0.05) |
| 17 | 0.23 (0.04) | 0.18 (0.04) | 0.20 (0.05) | 0.68 (0.04) | 0.57 (0.04) | 0.69 (0.06) | 0.34 (0.05) | 0.32 (0.07) | 0.28 (0.07) |
| 18 | 0.38 (0.08) | 0.31 (0.08) | 0.28 (0.10) | 1.42 (0.08) | 1.24 (0.09) | 1.43 (0.12) | 0.26 (0.05) | 0.25 (0.07) | 0.20 (0.07) |
| 19 | 0.09 (0.01) | 0.09 (0.02) | 0.08 (0..02) | 0.09 (0.01) | 0.09 (0.01) | 0.08 (0.02) | 1.00 (0.04) | 1.00 (0.06) | 1.00 (0.09) |
| 20 | 0.14 (0.02) | 0.07 (0.01) | 0.15 (0.04) | 0.22 (0.02) | 0.10 (0.01) | 0.32 (0.03) | 0.64 (0.06) | 0.63 (0.08) | 0.49 (0.10) |
| 21 | 0.02 (0.01) | 0.01 (0.00) | 0.02 (0.01) | 0.12 (0.01) | 0.10 (0.01) | 0.13 (0.01) | 0.15 (0.04) | 0.07 (0.04) | 0.18 (0.06) |
| 22 | 0.10 (0.01) | 0.07 (0.01) | 0.07 (0.01) | 0.10 (0.01) | 0.07 (0.01) | 0.08 (0.01) | 1.00 (0.01) | 1.00 (0.02) | 0.84 (0.05) |
| 23 | 0.01 (0.01) | 0.01 (0.01) | 0.00 (0.01) | 0.47 (0.03) | 0.35 (0.03) | 0.61 (0.05) | 0.01 (0.02) | 0.04 (0.03) | 0.00 (0.02) |
| 24 | 1.88 (0.22) | 0.51 (0.12) | 2.28 (0.34) | 2.71 (0.18) | 1.65 (0.13) | 2.94 (0.27) | 0.69 (0.05) | 0.31 (0.07) | 0.78 (0.07) |
| 25 | 2.13 (0.23) | 0.79 (0.15) | 2.63 (0.38) | 2.90 (0.19) | 1.60 (0.12) | 3.35 (0.30) | 0.74 (0.04) | 0.49 (0.07) | 0.78 (0.06) |
| 26 | 4.96 (0.39) | 0.33 (0.05) | 6.95 (0.81) | 5.18 (0.34) | 0.51 (0.04) | 7.42 (0.67) | 0.96 (0.02) | 0.65 (0.07) | 0.94 (0.04) |
| 27 | 0.01 (0.00) | 0.02 (0.00) | NA | 0.01 (0.00) | 0.02 (0.00) | NA | 0.91 (0.02) | 0.99 (0.03) | NA |
| 28 | 1.79 (0.00) | NA | NA | 1.79 (0.00) | NA | NA | 1.00 (0.00) | NA | NA |

**Table S3. GWAS results from spring and winter barley combined dataset**. For each GWAS peak, the quality check (QC) information is provided. QC class A means the peak is represented by two or more markers with significance above FDR of 0.05. QC class B means the peak is represented by one marker above FDR of 0.05 and two or more markers above FDR of 0.10. $r^2$ is not available for peaks that were detected in the second GWAS (similar to the initial GWAS but with the most significant marker as fixed effect). Since the marker data were coded as -1,0,1 where 1 is the minor allele in the initial marker data with 809 varieties, the effects here are defined with reference to minor allele. C: combined, S: spring, W: winter.

| Trait | GWAS peak | | | | | | | | | Minor allele frequency | | |
|---|---|---|---|---|---|---|---|---|---|---|---|---|
| | Marker | Chr | Pos (Mb) | Effect | $-\log_{10}p$ | $r^2$ | QC | Major | Minor | C | S | W |
| 1 | JHI-Hv50k-2016-250700 | 4H | 525069787 | 1.15 | 96.02 | 0.49 | A | G | T | 0.19 | 0.02 | 0.37 |
| 2 | JHI-Hv50k-2016-204917 | 3H | 631826807 | -0.49 | 6.40 | 0.07 | B | T | C | 0.49 | 0.05 | 0.98 |
| 3 | JHI-Hv50k-2016-36927 | 1H | 473266326 | -0.06 | 4.92 | NA | A | T | C | 0.16 | 0.01 | 0.32 |
| 3 | JHI-Hv50k-2016-269336 | 4H | 631675144 | 0.84 | 116.93 | 0.59 | A | G | C | 0.44 | 0.00 | 0.91 |
| 4 | JHI-Hv50k-2016-110539 | 2H | 676764470 | -1.01 | 42.77 | 0.28 | A | G | A | 0.25 | 0.00 | 0.53 |
| 4 | JHI-Hv50k-2016-468714 | 7H | 73547899 | -0.36 | 10.67 | NA | A | A | G | 0.10 | 0.02 | 0.19 |
| 8 | JHI-Hv50k-2016-110406 | 2H | 675755404 | -1.24 | 55.97 | 0.35 | A | A | G | 0.25 | 0.00 | 0.53 |
| 8 | JHI-Hv50k-2016-468714 | 7H | 73547899 | -0.37 | 11.01 | NA | A | A | G | 0.10 | 0.02 | 0.19 |
| 9 | JHI-Hv50k-2016-343 | 1H | 291116 | -0.37 | 6.23 | 0.04 | B | A | G | 0.07 | 0.07 | 0.06 |
| 9 | JHI-Hv50k-2016-62012 | 2H | 6179110 | -0.23 | 6.98 | 0.04 | B | T | A | 0.43 | 0.52 | 0.34 |
| 11 | SCRI_RS_168399 | 4H | 608432009 | -0.22 | 5.93 | 0.05 | A | A | C | 0.45 | 0.49 | 0.41 |
| 12 | JHI-Hv50k-2016-109271 | 2H | 663877140 | 0.16 | 6.05 | 0.05 | A | T | G | 0.30 | 0.24 | 0.37 |
| 12 | JHI-Hv50k-2016-326756 | 5H | 579729027 | -0.18 | 6.11 | 0.05 | B | G | A | 0.42 | 0.00 | 0.89 |
| 13 | JHI-Hv50k-2016-180055 | 3H | 437235827 | 0.65 | 5.56 | 0.03 | A | G | C | 0.05 | 0.00 | 0.11 |
| 19 | JHI-Hv50k-2016-107749 | 2H | 652418960 | -1.00 | 146.59 | 0.97 | A | G | A | 0.29 | 0.24 | 0.34 |
| 20 | JHI-Hv50k-2016-29039 | 1H | 404915672 | -0.60 | 11.39 | 0.09 | A | T | C | 0.30 | 0.02 | 0.60 |
| 20 | JHI-Hv50k-2016-108431 | 2H | 655814689 | -0.99 | 19.19 | 0.16 | A | G | A | 0.22 | 0.20 | 0.25 |
| 20 | JHI-Hv50k-2016-211623 | 3H | 659544025 | -0.29 | 4.12 | 0.03 | A | C | T | 0.16 | 0.01 | 0.32 |
| 20 | JHI-Hv50k-2016-308573 | 5H | 488462871 | -0.25 | 4.37 | NA | A | T | C | 0.06 | 0.02 | 0.11 |
| 21 | JHI-Hv50k-2016-461364 | 7H | 47563401 | 0.46 | 7.93 | 0.05 | A | A | C | 0.08 | 0.07 | 0.08 |
| 22 | JHI-Hv50k-2016-314851 | 5H | 542495869 | -0.38 | 24.72 | 0.15 | A | G | A | 0.36 | 0.52 | 0.18 |
| 23 | JHI-Hv50k-2016-498733 | 7H | 612520748 | -0.45 | 6.71 | 0.04 | A | C | T | 0.06 | 0.10 | 0.02 |
| 24 | JHI-Hv50k-2016-110449 | 2H | 676195259 | -0.70 | 44.08 | 0.27 | A | G | A | 0.34 | 0.14 | 0.56 |
| 24 | JHI-Hv50k-2016-468636 | 7H | 72974093 | -0.27 | 8.46 | NA | A | T | C | 0.22 | 0.03 | 0.43 |
| 25 | JHI-Hv50k-2016-104432 | 2H | 638369307 | 0.82 | 57.54 | 0.32 | A | A | G | 0.17 | 0.09 | 0.26 |
| 26 | JHI-Hv50k-2016-367876 | 6H | 330928 | 1.33 | 152.61 | 0.64 | A | G | A | 0.14 | 0.00 | 0.29 |
| 27 | JHI-Hv50k-2016-128079 | 2H | 724707499 | -1.00 | 24.45 | 0.16 | A | T | A | 0.48 | 0.05 | 0.95 |
| 28 | JHI-Hv50k-2016-41782 | 1H | 511924294 | -0.51 | 47.74 | 0.26 | A | C | G | 0.48 | 0.00 | 1.00 |
| 28 | JHI-Hv50k-2016-275542 | 4H | 643677857 | -0.63 | 63.95 | 0.34 | A | T | C | 0.48 | 0.00 | 1.00 |
| 28 | JHI-Hv50k-2016-323150 | 5H | 571032022 | -0.34 | 38.29 | 0.22 | A | G | A | 0.46 | 0.00 | 0.98 |

**Table S4. GWAS results from spring barley only dataset**. For each GWAS peak, the quality check (QC) information is provided. QC class A means the peak is represented by two or more markers with significance above FDR of 0.05. QC class B means the peak is represented by one marker above FDR of 0.05 and two or more markers above FDR of 0.10. Since the marker data were coded as -1,0,1 where 1 is the minor allele in the initial marker data with 809 varieties, the effects here are defined with reference to minor allele. C: combined, S: spring, W: winter.

| Trait | GWAS peak | | | | | | | | | Minor allele frequency | | |
|---|---|---|---|---|---|---|---|---|---|---|---|---|
| | Marker | Chr | Pos (Mb) | Effect | $-log_{10}p$ | $r^2$ | QC | Major | Minor | C | S | W |
| 1 | JHI-Hv50k-2016-250940 | 4H | 527125017 | 1.20 | 31.84 | 0.33 | A | G | T | 0.21 | 0.06 | 0.37 |
| 8 | JHI-Hv50k-2016-416214 | 6H | 536069651 | 0.51 | 6.75 | 0.12 | B | A | C | 0.30 | 0.16 | 0.45 |
| 9 | SCRI_RS_204276 | 1H | 248411 | -0.70 | 8.80 | 0.10 | A | C | T | 0.04 | 0.06 | 0.02 |
| 15 | JHI-Hv50k-2016-263040 | 4H | 608379093 | -0.25 | 4.76 | 0.07 | A | T | C | 0.45 | 0.49 | 0.41 |
| 19 | JHI-Hv50k-2016-107749 | 2H | 652418960 | -0.99 | 98.03 | 0.97 | A | G | A | 0.29 | 0.24 | 0.34 |
| 20 | JHI-Hv50k-2016-29941 | 1H | 414058556 | -0.57 | 4.53 | 0.07 | A | C | G | 0.31 | 0.06 | 0.58 |
| 20 | JHI-Hv50k-2016-108561 | 2H | 656596131 | -0.75 | 8.01 | 0.12 | A | C | T | 0.23 | 0.22 | 0.25 |
| 22 | JHI-Hv50k-2016-316002 | 5H | 546260097 | -1.04 | 194.19 | 0.91 | A | G | C | 0.22 | 0.34 | 0.10 |
| 23 | JHI-Hv50k-2016-498708 | 7H | 612518575 | -0.43 | 4.55 | 0.05 | A | T | C | 0.05 | 0.10 | 0.00 |
| 24 | JHI-Hv50k-2016-110405 | 2H | 675754697 | -0.68 | 19.42 | 0.25 | A | A | G | 0.36 | 0.18 | 0.57 |
| 25 | JHI-Hv50k-2016-104388 | 2H | 638193925 | 1.31 | 44.49 | 0.43 | A | C | T | 0.05 | 0.09 | 0.00 |
| 27 | JHI-Hv50k-2016-128079 | 2H | 724707499 | -1.31 | 27.93 | 0.30 | A | T | A | 0.48 | 0.05 | 0.95 |

**Table S5. GWAS results from winter barley only dataset**. For each GWAS peak, the quality check (QC) information is provided. QC class A means the peak is represented by two or more markers with significance above FDR of 0.05. QC class B means the peak is represented by one marker above FDR of 0.05 and two or more markers above FDR of 0.10. $r^2$ is not available for peaks that were detected in second GWAS (similar to the initial GWAS but with the most significant marker as fixed effect). Since the marker data were coded as -1,0,1 where 1 is the minor allele in the initial marker data with 809 varieties, the effects here are defined with reference to minor allele. C: combined, S: spring, W: winter.

| Trait | GWAS peak | | | | | | | | | Minor allele frequency | | |
|---|---|---|---|---|---|---|---|---|---|---|---|---|
| | Marker | Chr | Pos (Mb) | Effect | $-\log_{10}p$ | $r^2$ | QC | Major | Minor | C | S | W |
| 1 | JHI-Hv50k-2016-251355 | 4H | 531530594 | 1.00 | 59.89 | 0.59 | A | T | C | 0.32 | 0.27 | 0.37 |
| 3 | JHI-Hv50k-2016-269297 | 4H | 631635187 | 1.49 | 61.39 | 0.59 | A | G | C | 0.44 | 0.00 | 0.91 |
| 4 | JHI-Hv50k-2016-110120 | 2H | 674233412 | -0.78 | 26.64 | 0.43 | A | A | G | 0.26 | 0.02 | 0.53 |
| 8 | JHI-Hv50k-2016-110120 | 2H | 674233412 | -0.94 | 40.39 | 0.56 | A | A | G | 0.26 | 0.02 | 0.53 |
| 8 | JHI-Hv50k-2016-468636 | 7H | 72974093 | -0.18 | 5.42 | NA | A | T | C | 0.22 | 0.03 | 0.43 |
| 9 | JHI-Hv50k-2016-343 | 1H | 291116 | -0.82 | 10.49 | 0.13 | A | A | G | 0.07 | 0.07 | 0.06 |
| 12 | JHI-Hv50k-2016-109317 | 2H | 664191672 | 0.18 | 5.52 | 0.07 | A | A | T | 0.31 | 0.25 | 0.37 |
| 19 | JHI-Hv50k-2016-108296 | 2H | 655509717 | -0.99 | 21.40 | 0.80 | A | G | T | 0.24 | 0.24 | 0.25 |
| 20 | JHI-Hv50k-2016-29031 | 1H | 404914383 | -0.55 | 7.33 | 0.13 | A | A | G | 0.30 | 0.02 | 0.60 |
| 20 | JHI-Hv50k-2016-107816 | 2H | 653415012 | -0.87 | 9.20 | 0.17 | A | T | C | 0.30 | 0.24 | 0.37 |
| 21 | JHI-Hv50k-2016-460614 | 7H | 42684996 | 1.29 | 14.60 | 0.22 | A | A | T | 0.08 | 0.10 | 0.06 |
| 22 | JHI-Hv50k-2016-316000 | 5H | 546259246 | -1.43 | 64.16 | 0.59 | A | C | G | 0.22 | 0.34 | 0.10 |
| 24 | JHI-Hv50k-2016-110120 | 2H | 674233412 | -0.86 | 33.11 | 0.39 | A | A | G | 0.26 | 0.02 | 0.53 |
| 24 | JHI-Hv50k-2016-468636 | 7H | 72974093 | -0.23 | 7.44 | NA | A | T | C | 0.22 | 0.03 | 0.43 |
| 25 | JHI-Hv50k-2016-105043 | 2H | 640060225 | 0.84 | 26.69 | 0.38 | A | T | A | 0.14 | 0.03 | 0.26 |
| 26 | JHI-Hv50k-2016-368372 | 6H | 1005358 | 1.04 | 102.85 | 0.79 | A | A | C | 0.20 | 0.10 | 0.30 |

**Table S6. Comparison of simulated progeny with small marker set**. For each variety, we compared the markers from the simulated progeny to its real sibling (variety), parents and among themselves. Note: N-seg is the number of segregating markers between each parent pairs, N-unique is the number of unique haplotypes found in the simulated progeny, and Non-unique is the percentage of simulated progeny with over 1% probability of matching with other simulated progeny.

| Variety | AFP | Season | N-seg | N-unique | Sim vs Variety (%) | Sim vs Parent 1 (%) | Sim vs Parent 2 (%) | Non-unique (%) |
|---|---|---|---|---|---|---|---|---|
| SW Alison | 1788 | alt | 15 | 6238 | 0 | 0.06 | 0.09 | 0 |
| Natasha | 545 | spring | 15 | 6721 | 0 | 0.03 | 0.09 | 0 |
| Franklin | 1200 | spring | 14 | 6639 | 0.01 | 0.08 | 0.09 | 0 |
| Wren | 1242 | spring | 17 | 7806 | 0 | 0.06 | 0.03 | 0 |
| Toddy | 1298 | spring | 14 | 7251 | 0 | 0.03 | 0.03 | 0 |
| Decanter | 1411 | spring | 16 | 7804 | 0.04 | 0.02 | 0.03 | 0 |
| Horizon | 1423 | spring | 13 | 4299 | 0 | 0.12 | 0.22 | 0 |
| Static | 1427 | spring | 15 | 4407 | 0 | 0.12 | 0.19 | 0 |
| Berwick | 1474 | spring | 14 | 5064 | 0 | 0.12 | 0.11 | 0 |
| Thrift | 1554 | spring | 16 | 6821 | 0.01 | 0.11 | 0.06 | 0 |
| Akita | 1555 | spring | 12 | 5514 | 0 | 0.08 | 0.07 | 0 |
| Dew | 1556 | spring | 19 | 8499 | 0 | 0.04 | 0.03 | 0 |
| Alliot | 1563 | spring | 15 | 6612 | 0.02 | 0.07 | 0.12 | 0 |
| Prestige | 1566 | spring | 14 | 4620 | 0 | 0.11 | 0.11 | 0 |
| Protege | 1567 | spring | 14 | 5122 | 0 | 0.09 | 0.06 | 0 |
| Amphora | 1588 | spring | 14 | 6487 | 0 | 0.1 | 0.08 | 0 |
| Agenda | 1589 | spring | 12 | 3190 | 0.04 | 0.23 | 0.21 | 0 |
| Azure | 1591 | spring | 16 | 6410 | 0 | 0.15 | 0.12 | 0 |
| Sacha | 1651 | spring | 19 | 8506 | 0 | 0.04 | 0.01 | 0 |
| NSL 98-5065 | 1663 | spring | 12 | 3998 | 0 | 0.24 | 0.25 | 0 |
| Adonis | 1666 | spring | 18 | 8777 | 0 | 0.02 | 0.04 | 0 |
| Spire | 1668 | spring | 12 | 4712 | 0.04 | 0.11 | 0.11 | 0 |
| Spike | 1669 | spring | 12 | 4761 | 0.07 | 0.17 | 0.05 | 0 |
| Catalina | 1746 | spring | 14 | 4288 | 0.01 | 0.18 | 0.16 | 0 |
| Global | 1761 | spring | 13 | 5702 | 0.01 | 0.03 | 0.07 | 0 |
| Maypole | 1775 | spring | 15 | 5451 | 0 | 0.09 | 0.18 | 0 |
| Class | 1777 | spring | 12 | 4449 | 0.01 | 0.12 | 0.12 | 0 |
| Brazil | 1780 | spring | 13 | 3680 | 0.05 | 0.22 | 0.14 | 0 |
| Macarena | 1837 | spring | 10 | 3101 | 0.18 | 0.2 | 0.17 | 0 |
| Beryllium | 1847 | spring | 10 | 2708 | 0.07 | 0.21 | 0.18 | 0 |
| Toby | 1864 | spring | 12 | 3607 | 0.05 | 0.35 | 0.26 | 0 |
| Doyen | 1865 | spring | 13 | 6463 | 0.02 | 0.1 | 0.03 | 0 |
| Oxbridge | 1940 | spring | 21 | 9407 | 0 | 0.03 | 0 | 0 |
| CPBT B67 | 1947 | spring | 11 | 4202 | 0.03 | 0.11 | 0.09 | 0 |
| Macaw | 1955 | spring | 12 | 2787 | 0.43 | 0.27 | 0.39 | 0 |
| Wicket | 1968 | spring | 12 | 3186 | 0.06 | 0.22 | 0.24 | 0 |
| Polanski | 2028 | spring | 10 | 1864 | 0.56 | 0.61 | 0.6 | 0 |
| Rummy | 2034 | spring | 12 | 2530 | 0.15 | 0.48 | 0.46 | 0 |
| Cribbage | 2039 | spring | 13 | 5761 | 0 | 0.09 | 0.07 | 0 |
| Silicon | 2040 | spring | 18 | 8800 | 0.01 | 0.01 | 0.02 | 0 |
| 995964 | 2044 | spring | 12 | 3827 | 0.15 | 0.19 | 0.11 | 0 |
| Tucson | 2050 | spring | 13 | 5319 | 0 | 0.1 | 0.18 | 0 |
| Putney | 2053 | spring | 15 | 6926 | 0 | 0.08 | 0.07 | 0 |
| Centurion | 2054 | spring | 14 | 4660 | 0 | 0.09 | 0.11 | 0 |
| Scandium | 2096 | spring | 12 | 3340 | 0 | 0.21 | 0.22 | 0 |
| Token | 2103 | spring | 7 | 691 | 1.63 | 1.78 | 1.73 | 52.09 |
| Calico | 2104 | spring | 12 | 3919 | 0.02 | 0.24 | 0.22 | 0 |
| Shakira | 2110 | spring | 10 | 2835 | 0 | 0.29 | 0.34 | 0 |

| | | | | | | | | |
|---|---|---|---|---|---|---|---|---|
| Publican | 2119 | spring | 17 | 5766 | 0.03 | 0.18 | 0.12 | 0 |
| Spotlight | 2120 | spring | 12 | 2481 | 0 | 0.54 | 0.6 | 0 |
| Quench | 2121 | spring | 17 | 5758 | 0.01 | 0.15 | 0.15 | 0 |
| Taphouse | 2122 | spring | 11 | 1584 | 0.56 | 0.86 | 0.83 | 0 |
| Balalaika | 2173 | spring | 17 | 8087 | 0 | 0.03 | 0.04 | 0 |
| Azalea | 2180 | spring | 11 | 1885 | 0.02 | 0.7 | 0.59 | 0 |
| Ingmar | 2187 | spring | 17 | 7911 | 0.03 | 0.04 | 0.03 | 0 |
| Tamise | 2188 | spring | 11 | 4320 | 0 | 0.1 | 0.15 | 0 |
| Belgravia | 2201 | spring | 11 | 2825 | 0.08 | 0.75 | 0.51 | 0 |
| Knightsbridge | 2203 | spring | 14 | 4532 | 0.19 | 0.24 | 0.15 | 0 |
| Sweeney | 2206 | spring | 16 | 5514 | 0.02 | 0.16 | 0.14 | 0 |
| Snakebite | 2207 | spring | 11 | 1615 | 0.02 | 0.78 | 0.91 | 0 |
| Maltby | 2209 | spring | 12 | 4267 | 0.03 | 0.14 | 0.15 | 0 |
| Jolika | 2212 | spring | 17 | 5717 | 0.02 | 0.1 | 0.15 | 0 |
| Twister | 2272 | spring | 14 | 4572 | 0.01 | 0.14 | 0.16 | 0 |
| Cropton | 2276 | spring | 16 | 7325 | 0 | 0.07 | 0.02 | 0 |
| Concerto | 2288 | spring | 11 | 2787 | 0.04 | 0.69 | 0.68 | 0 |
| Virgil | 2289 | spring | 11 | 2848 | 0.01 | 0.55 | 0.54 | 0 |
| Garner | 2335 | spring | 11 | 2121 | 0.19 | 0.68 | 0.83 | 0 |
| Propino | 2336 | spring | 16 | 5703 | 0 | 0.22 | 0.19 | 0 |
| Cairn | 2338 | spring | 12 | 2992 | 0.23 | 0.4 | 0.32 | 0 |
| Checkmate | 2396 | spring | 13 | 4249 | 0.02 | 0.16 | 0.15 | 0 |
| SY Taberna | 2398 | spring | 11 | 2471 | 0.37 | 0.3 | 0.48 | 0 |
| Muirhead | 2401 | spring | 14 | 5301 | 0.02 | 0.11 | 0.12 | 0 |
| Summit | 2402 | spring | 11 | 2105 | 0.09 | 0.77 | 0.62 | 0 |
| Panther | 2418 | spring | 17 | 8314 | 0 | 0.04 | 0.04 | 0 |
| Western | 2430 | spring | 14 | 5581 | 0 | 0.16 | 0.09 | 0 |
| Overture | 2465 | spring | 15 | 5971 | 0 | 0.05 | 0.11 | 0 |
| Chronicle | 2466 | spring | 15 | 6075 | 0 | 0.07 | 0.12 | 0 |
| Odyssey | 2470 | spring | 15 | 6017 | 0 | 0.07 | 0.07 | 0 |
| Shandy | 2478 | spring | 8 | 1143 | 0.11 | 1.6 | 1.62 | 26.35 |
| Columbus | 2481 | spring | 15 | 4992 | 0.15 | 0.11 | 0.1 | 0 |
| SY Barrell | 2497 | spring | 17 | 7258 | 0 | 0.1 | 0.03 | 0 |
| Tesla | 2541 | spring | 15 | 5372 | 0.02 | 0.15 | 0.17 | 0 |
| Momentum | 2542 | spring | 15 | 6097 | 0 | 0.08 | 0.14 | 0 |
| Convivial | 2557 | spring | 19 | 8769 | 0 | 0.03 | 0.03 | 0 |
| Crooner | 2560 | spring | 14 | 5514 | 0.02 | 0.07 | 0.05 | 0 |
| Mickle | 2567 | spring | 9 | 1654 | 0.65 | 0.65 | 0.66 | 0 |
| Glassel | 2570 | spring | 16 | 7160 | 0 | 0.05 | 0.03 | 0 |
| Sanette | 2572 | spring | 10 | 1762 | 0.02 | 0.79 | 0.8 | 0 |
| KWS Irina | 2613 | spring | 9 | 860 | 0.02 | 1.31 | 1.51 | 48.38 |
| Acumen | 2623 | spring | 6 | 214 | 4.06 | 4.05 | 4.57 | 84.29 |
| Malt Jagger | 2626 | spring | 14 | 5389 | 0 | 0.13 | 0.1 | 0 |
| Hacker | 2627 | spring | 16 | 6948 | 0 | 0.08 | 0.1 | 0 |
| Shada | 2642 | spring | 7 | 453 | 2.54 | 2.43 | 2.23 | 62.83 |
| Shaloo | 2646 | spring | 13 | 2934 | 0.02 | 0.32 | 0.48 | 0 |
| Sienna | 2697 | spring | 10 | 2256 | 0.07 | 0.27 | 0.34 | 0 |
| Octavia | 2699 | spring | 11 | 1907 | 0 | 1.02 | 0.93 | 9.38 |
| Olympus | 2701 | spring | 10 | 1486 | 0.17 | 0.77 | 0.73 | 0 |
| Deveron | 2702 | spring | 10 | 1516 | 0.81 | 0.81 | 0.84 | 3.16 |
| KWS Sassy | 2771 | spring | 14 | 5027 | 0 | 0.16 | 0.14 | 0 |
| Laureate | 2780 | spring | 13 | 5001 | 0.08 | 0.13 | 0.19 | 0 |
| Ovation | 2788 | spring | 7 | 501 | 0.05 | 3.33 | 3.2 | 62.39 |
| Orbital | 2790 | spring | 9 | 1257 | 0.05 | 1.03 | 0.97 | 32.44 |
| Acorn | 2838 | spring | 13 | 4288 | 0.01 | 0.16 | 0.22 | 0 |
| Dioptric | 2870 | spring | 8 | 1330 | 0.34 | 0.59 | 0.78 | 0 |
| LG Mojito | 2902 | spring | 12 | 2917 | 0.03 | 0.39 | 0.41 | 0 |
| LG Figaro | 2903 | spring | 10 | 3150 | 0.05 | 0.17 | 0.25 | 0 |
| LG Diablo | 2907 | spring | 10 | 3085 | 0.01 | 0.23 | 0.19 | 0 |

| | | | | | | | | |
|---|---|---|---|---|---|---|---|---|
| RGT Orbiter | 2979 | spring | 14 | 3381 | 0.33 | 0.31 | 0.43 | 0 |
| Accurance | 2986 | spring | 11 | 1393 | 1.65 | 1.6 | 1.7 | 27.16 |
| KWS Elsie | 2993 | spring | 8 | 436 | 1.38 | 3.24 | 3.81 | 76.47 |
| Gulliver | 2995 | spring | 9 | 1338 | 0.07 | 1.16 | 1.08 | 17.95 |
| LG Goddess | 2999 | spring | 4 | 72 | 7.51 | 7.97 | 7.83 | 88.4 |
| Embrace | 3004 | spring | 13 | 3983 | 0 | 0.42 | 0.37 | 0 |
| SY Dolomite | 3016 | spring | 9 | 824 | 2.43 | 2.34 | 2.26 | 46.73 |
| SY Kailash | 3017 | spring | 9 | 749 | 2.17 | 2.07 | 2.74 | 49.33 |
| Gypsy | 884 | winter | 19 | 7848 | 0 | 0.05 | 0.04 | 0 |
| Duet | 1161 | winter | 12 | 2631 | 0.1 | 0.43 | 0.4 | 0 |
| Sunrise | 1162 | winter | 21 | 8719 | 0 | 0.02 | 0.02 | 0 |
| Madrigal | 1172 | winter | 16 | 6738 | 0 | 0.04 | 0.01 | 0 |
| Medoc | 1212 | winter | 16 | 6690 | 0 | 0.09 | 0.04 | 0 |
| Muscat | 1214 | winter | 8 | 1270 | 0.65 | 0.72 | 0.93 | 0 |
| Gleam | 1228 | winter | 21 | 8749 | 0 | 0.01 | 0.06 | 0 |
| Falcon | 1260 | winter | 13 | 4706 | 0.03 | 0.09 | 0.11 | 0 |
| Glint | 1271 | winter | 21 | 8721 | 0 | 0.05 | 0.02 | 0 |
| Oriflamme | 1272 | winter | 17 | 6608 | 0 | 0.09 | 0.06 | 0 |
| Antigua | 1276 | winter | 16 | 6765 | 0 | 0.07 | 0.06 | 0 |
| Spice | 1285 | winter | 18 | 8697 | 0 | 0.03 | 0.02 | 0 |
| Ravel | 1317 | winter | 17 | 6578 | 0.01 | 0.04 | 0.1 | 0 |
| Pearl | 1318 | winter | 18 | 5298 | 0 | 0.19 | 0.15 | 0 |
| Jewel | 1320 | winter | 12 | 3032 | 0.3 | 0.32 | 0.28 | 0 |
| Peridot | 1321 | winter | 19 | 8439 | 0 | 0.03 | 0.03 | 0 |
| Baton | 1326 | winter | 18 | 8101 | 0 | 0.02 | 0.01 | 0 |
| Goldrush | 1381 | winter | 18 | 8644 | 0 | 0.01 | 0.02 | 0 |
| Becket | 1385 | winter | 15 | 5993 | 0.03 | 0.09 | 0.09 | 0 |
| Chord | 1387 | winter | 15 | 6060 | 0 | 0.07 | 0.1 | 0 |
| Mahogany | 1388 | winter | 14 | 4634 | 0.07 | 0.07 | 0.05 | 0 |
| Magnolia | 1390 | winter | 18 | 8079 | 0 | 0.01 | 0.05 | 0 |
| Bistro | 1392 | winter | 13 | 4562 | 0 | 0.05 | 0.04 | 0 |
| Heligan | 1402 | winter | 16 | 7066 | 0 | 0.02 | 0.02 | 0 |
| Tabetha | 1404 | winter | 17 | 7135 | 0 | 0.07 | 0.09 | 0 |
| Molly | 1405 | winter | 16 | 6874 | 0 | 0.05 | 0.06 | 0 |
| Hurricane | 1441 | winter | 16 | 6913 | 0 | 0.06 | 0.1 | 0 |
| Wizard | 1442 | winter | 16 | 6913 | 0.01 | 0.1 | 0.04 | 0 |
| Arrow | 1445 | winter | 15 | 4860 | 0.07 | 0.15 | 0.15 | 0 |
| Anvil | 1511 | winter | 12 | 4202 | 0.04 | 0.21 | 0.1 | 0 |
| Haka | 1528 | winter | 16 | 6926 | 0 | 0.02 | 0.02 | 0 |
| CPBT B28 | 1529 | winter | 18 | 7344 | 0 | 0.03 | 0.06 | 0 |
| Harland | 1530 | winter | 18 | 7328 | 0 | 0.05 | 0.02 | 0 |
| Campion | 1531 | winter | 17 | 7155 | 0.04 | 0.01 | 0.06 | 0 |
| Chamomile | 1532 | winter | 17 | 7377 | 0.04 | 0.06 | 0.08 | 0 |
| Montage | 1541 | winter | 10 | 1967 | 0.11 | 0.91 | 0.85 | 0 |
| Sumo | 1547 | winter | 16 | 6924 | 0 | 0.06 | 0.04 | 0 |
| Barcelona | 1552 | winter | 16 | 6285 | 0 | 0.08 | 0.01 | 0 |
| Sapphire | 1605 | winter | 8 | 382 | 2.45 | 2.64 | 2.53 | 78.86 |
| Tucker | 1608 | winter | 10 | 1943 | 0.07 | 0.78 | 0.66 | 0 |
| Tipster | 1610 | winter | 8 | 725 | 0.02 | 3.34 | 3.39 | 63.43 |
| Rafiki | 1618 | winter | 11 | 1527 | 0.51 | 1.46 | 1.2 | 21.79 |
| Scylla | 1628 | winter | 18 | 7448 | 0 | 0.07 | 0.04 | 0 |
| Caption | 1630 | winter | 17 | 7305 | 0 | 0.06 | 0.11 | 0 |
| Jessica | 1693 | winter | 16 | 5357 | 0.08 | 0.14 | 0.13 | 0 |
| Cannock | 1700 | winter | 15 | 4078 | 0 | 0.21 | 0.33 | 0 |
| NSL 97-6002 | 1701 | winter | 16 | 5064 | 0 | 0.14 | 0.16 | 0 |
| Mead | 1728 | winter | 11 | 1888 | 0.33 | 0.66 | 0.78 | 0 |
| SW Farrier | 1733 | winter | 15 | 3682 | 0.04 | 0.19 | 0.17 | 0 |
| Louise | 1783 | winter | 16 | 5466 | 0 | 0.11 | 0.2 | 0 |
| SW Sienna | 1787 | winter | 18 | 7871 | 0 | 0.05 | 0.07 | 0 |

| | | | | | | | | |
|---|---|---|---|---|---|---|---|---|
| Nocturne | 1792 | winter | 16 | 6753 | 0 | 0.07 | 0.13 | 0 |
| Wombat | 1793 | winter | 11 | 2566 | 0.13 | 0.3 | 0.24 | 0 |
| Kingston | 1799 | winter | 8 | 1310 | 0.11 | 0.53 | 0.6 | 0 |
| Lambada | 1808 | winter | 13 | 2066 | 0 | 0.71 | 0.63 | 0 |
| Mortimer | 1824 | winter | 16 | 5368 | 0.04 | 0.2 | 0.21 | 0 |
| Saffron | 1880 | winter | 13 | 4066 | 0.14 | 0.14 | 0.18 | 0 |
| CPBT B66 | 1882 | winter | 22 | 9039 | 0 | 0.01 | 0.04 | 0 |
| Amarena | 1886 | winter | 7 | 416 | 3.16 | 2.96 | 3.34 | 79.2 |
| Faraday | 1892 | winter | 15 | 5772 | 0 | 0.08 | 0.07 | 0 |
| Coriolis | 1894 | winter | 15 | 4273 | 0.15 | 0.22 | 0.26 | 0 |
| 961374 | 1900 | winter | 18 | 6051 | 0.02 | 0.08 | 0.09 | 0 |
| Rattle | 1914 | winter | 9 | 1348 | 0.59 | 0.43 | 0.39 | 0 |
| Celebrity | 1982 | winter | 16 | 4490 | 0 | 0.18 | 0.15 | 0 |
| Cypress | 1983 | winter | 12 | 2738 | 0.04 | 0.34 | 0.37 | 0 |
| Cinnamon | 1984 | winter | 20 | 8595 | 0 | 0.02 | 0.02 | 0 |
| Archimedes | 1989 | winter | 16 | 6317 | 0 | 0.11 | 0.04 | 0 |
| Celsius | 1990 | winter | 12 | 3179 | 0 | 0.16 | 0.27 | 0 |
| Fahrenheit | 1991 | winter | 12 | 3220 | 0.07 | 0.14 | 0.22 | 0 |
| Silverstone | 2017 | winter | 14 | 4302 | 0.09 | 0.23 | 0.18 | 0 |
| Suzuka | 2083 | winter | 10 | 2123 | 0.16 | 0.4 | 0.33 | 0 |
| Cedar | 2088 | winter | 12 | 3684 | 0 | 0.14 | 0.2 | 0 |
| CPBT B78 | 2134 | winter | 8 | 1372 | 0.21 | 0.43 | 0.56 | 0 |
| 410/3E | 2145 | winter | 17 | 5077 | 0 | 0.11 | 0.21 | 0 |
| AC 99/077/2 | 2160 | winter | 17 | 6723 | 0 | 0.06 | 0.07 | 0 |
| Salling | 2228 | winter | 7 | 664 | 1.11 | 1.12 | 1.14 | 59.69 |
| Metaxa | 2237 | winter | 17 | 6727 | 0.02 | 0.05 | 0.05 | 0 |
| Blazing | 2375 | winter | 12 | 3040 | 0.08 | 0.33 | 0.32 | 0 |
| KWS Salsa | 2433 | winter | 15 | 6630 | 0 | 0.04 | 0 | 0 |
| SY Ninkasi | 2442 | winter | 15 | 3835 | 0.17 | 0.36 | 0.34 | 0 |
| Talisman | 2517 | winter | 13 | 3371 | 0.12 | 0.18 | 0.21 | 0 |
| KWS Glacier | 2523 | winter | 13 | 4254 | 0.07 | 0.16 | 0.23 | 0 |
| Cadillac | 2581 | winter | 16 | 8101 | 0.03 | 0.02 | 0.03 | 0 |
| Cavalier | 2582 | winter | 16 | 8058 | 0 | 0.03 | 0.01 | 0 |
| Harlequin | 2583 | winter | 16 | 8100 | 0.03 | 0.02 | 0.02 | 0 |
| Sandra | 2585 | winter | 16 | 6721 | 0 | 0.03 | 0.05 | 0 |
| Acute | 2586 | winter | 9 | 780 | 0.36 | 1.1 | 1.12 | 30.76 |
| Tetris | 2602 | winter | 12 | 3619 | 0.03 | 0.1 | 0.1 | 0 |
| KWS Infinity | 2650 | winter | 13 | 4244 | 0.06 | 0.14 | 0.13 | 0 |
| Perseus | 2651 | winter | 16 | 8020 | 0.01 | 0.05 | 0.01 | 0 |
| KWS Tower | 2725 | winter | 12 | 3114 | 0.1 | 0.43 | 0.22 | 0 |
| KWS Orwell | 2728 | winter | 13 | 3136 | 0 | 0.28 | 0.21 | 0 |
| KWS Carbis | 2802 | winter | 14 | 4400 | 0.04 | 0.15 | 0.06 | 0 |
| KWS Creswell | 2804 | winter | 14 | 4412 | 0.03 | 0.16 | 0.17 | 0 |
| Coref | 2887 | winter | 17 | 5447 | 0.02 | 0.21 | 0.2 | 0 |
| Adelac | 2956 | winter | 13 | 4601 | 0.11 | 0.13 | 0.13 | 0 |
| LG Mountain | 2976 | winter | 8 | 1278 | 0.14 | 0.65 | 0.77 | 0 |

**Table S7. Summary of Manhattan distances from varying number of random markers**.

| Number of marker | Manhattan distance | |
|---|---|---|
| | Mean | Variance |
| 1 | 0.912 | 0.948 |
| 2 | 0.951 | 0.629 |
| 3 | 0.465 | 0.251 |
| 4 | 0.727 | 0.264 |
| 5 | 0.813 | 0.205 |
| 6 | 0.758 | 0.156 |
| 8 | 0.673 | 0.104 |
| 10 | 0.804 | 0.110 |
| 13 | 0.510 | 0.077 |
| 16 | 0.595 | 0.066 |
| 20 | 0.672 | 0.094 |
| 25 | 0.472 | 0.032 |
| 32 | 0.514 | 0.031 |
| 40 | 0.615 | 0.045 |
| 50 | 0.529 | 0.027 |
| 63 | 0.608 | 0.033 |
| 79 | 0.588 | 0.030 |
| 100 | 0.583 | 0.024 |
| 126 | 0.555 | 0.023 |
| 158 | 0.618 | 0.025 |
| 200 | 0.593 | 0.020 |
| 251 | 0.637 | 0.031 |
| 316 | 0.609 | 0.024 |
| 398 | 0.580 | 0.026 |
| 501 | 0.607 | 0.023 |
| 631 | 0.603 | 0.020 |
| 794 | 0.605 | 0.023 |
| 1000 | 0.606 | 0.021 |
| 1259 | 0.604 | 0.025 |
| 1585 | 0.611 | 0.023 |
| 1995 | 0.601 | 0.020 |
| 2512 | 0.609 | 0.021 |
| 3162 | 0.605 | 0.022 |
| 3981 | 0.612 | 0.023 |
| 5012 | 0.611 | 0.022 |
| 6310 | 0.603 | 0.023 |
| 7943 | 0.602 | 0.022 |
| 10000 | 0.603 | 0.022 |
| 12589 | 0.605 | 0.021 |
| 15849 | 0.605 | 0.022 |
| 19953 | 0.602 | 0.021 |
| 25119 | 0.605 | 0.022 |
| 31623 | 0.603 | 0.022 |
| 40065 | 0.604 | 0.022 |

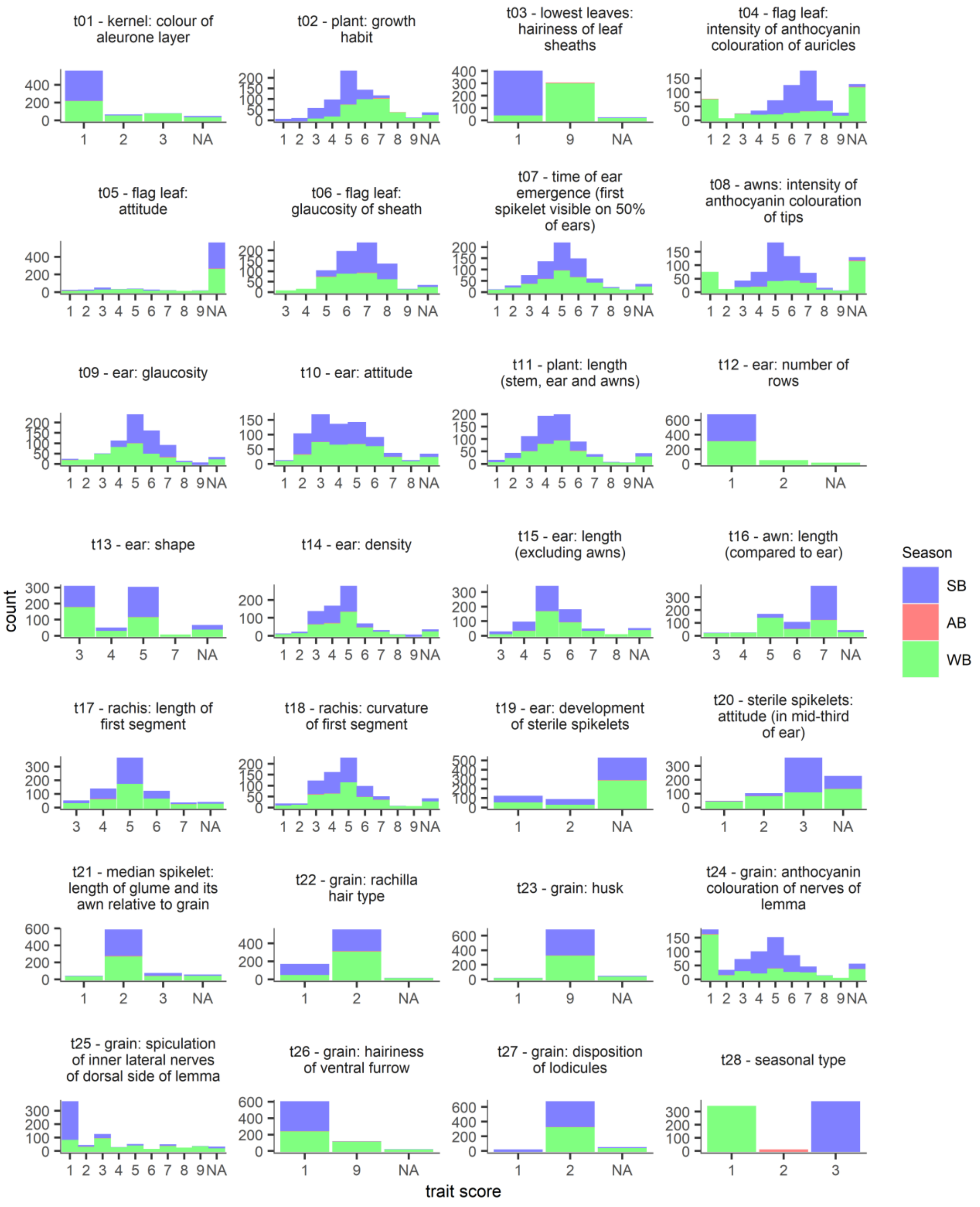


**Figure S1. Distributions of DUS trait scores.** Within each DUS trait, the bars are filled according to seasonal type. SB: spring barley, AB: alternative barley, WB: winter barley.

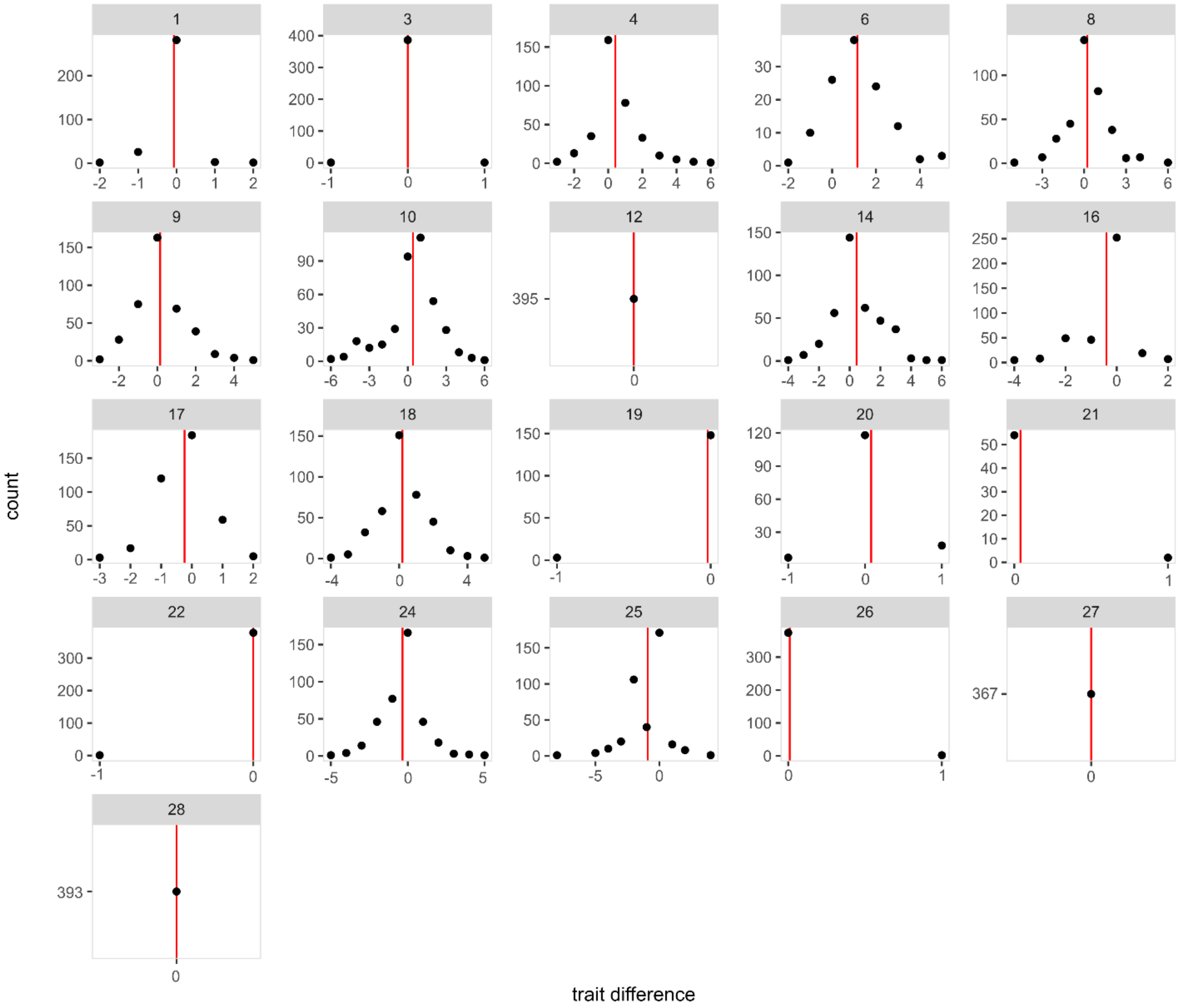


**Figure S2. Distributions of DUS trait score differences between NIAB and SASA.** DUS trait score differences are calculated as trait (NIAB) - trait (SASA). Negative differences mean that NIAB trait score is lower than SASA trait score, and vice versa. There is no variety in common between the NIAB and SASA datasets for 7 traits, and thus no comparison is available for these traits. The trait names are shown above each plot and the means of differences are shown as vertical red lines.

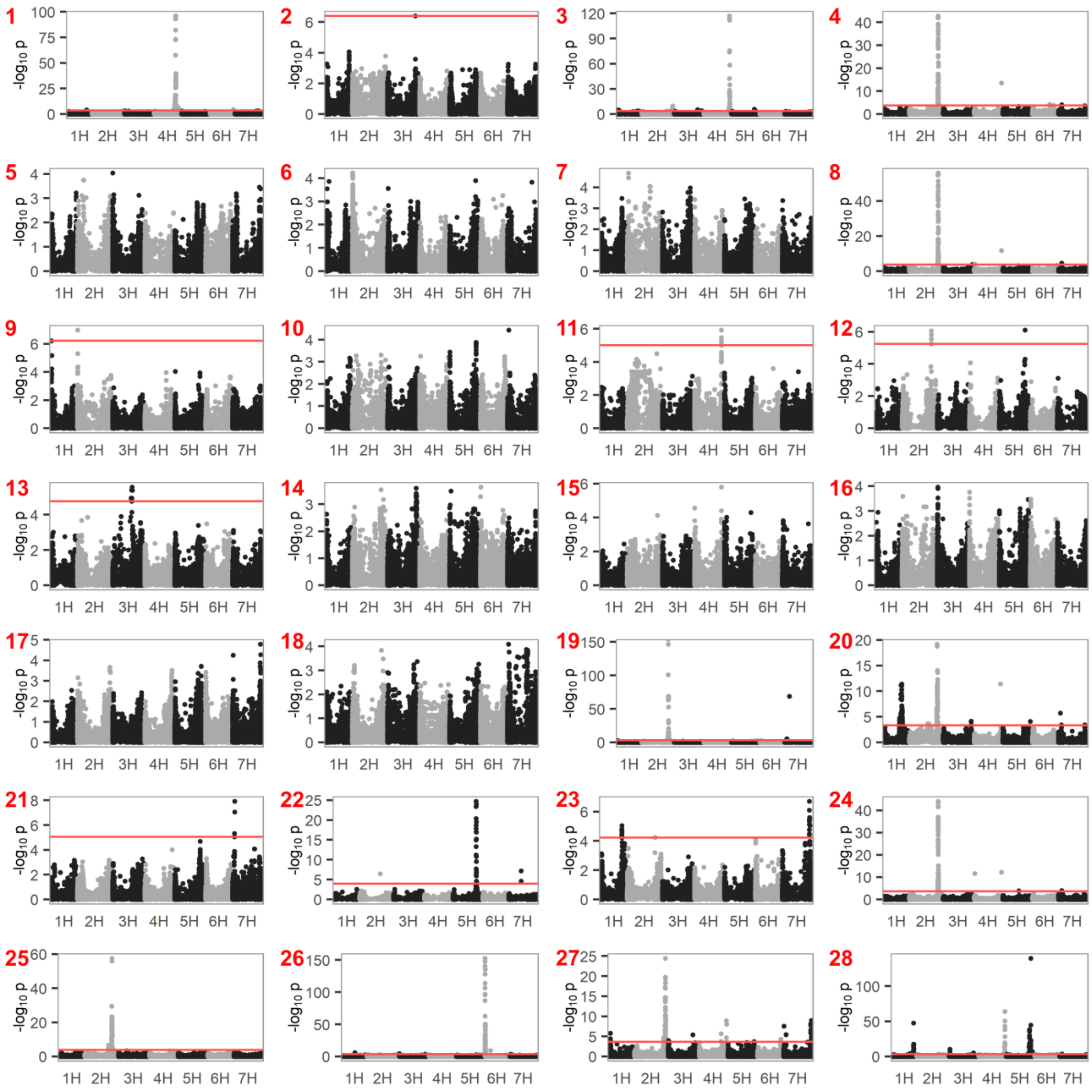

**Fig. S3. Manhattan plots of GWAS results from spring and winter barley combined datasets (n=710)**. Trait numbers are annotated at the top left corner of each Manhattan plot. Red horizontal line represents FDR = 0.05. If absent no significant marker was identified.

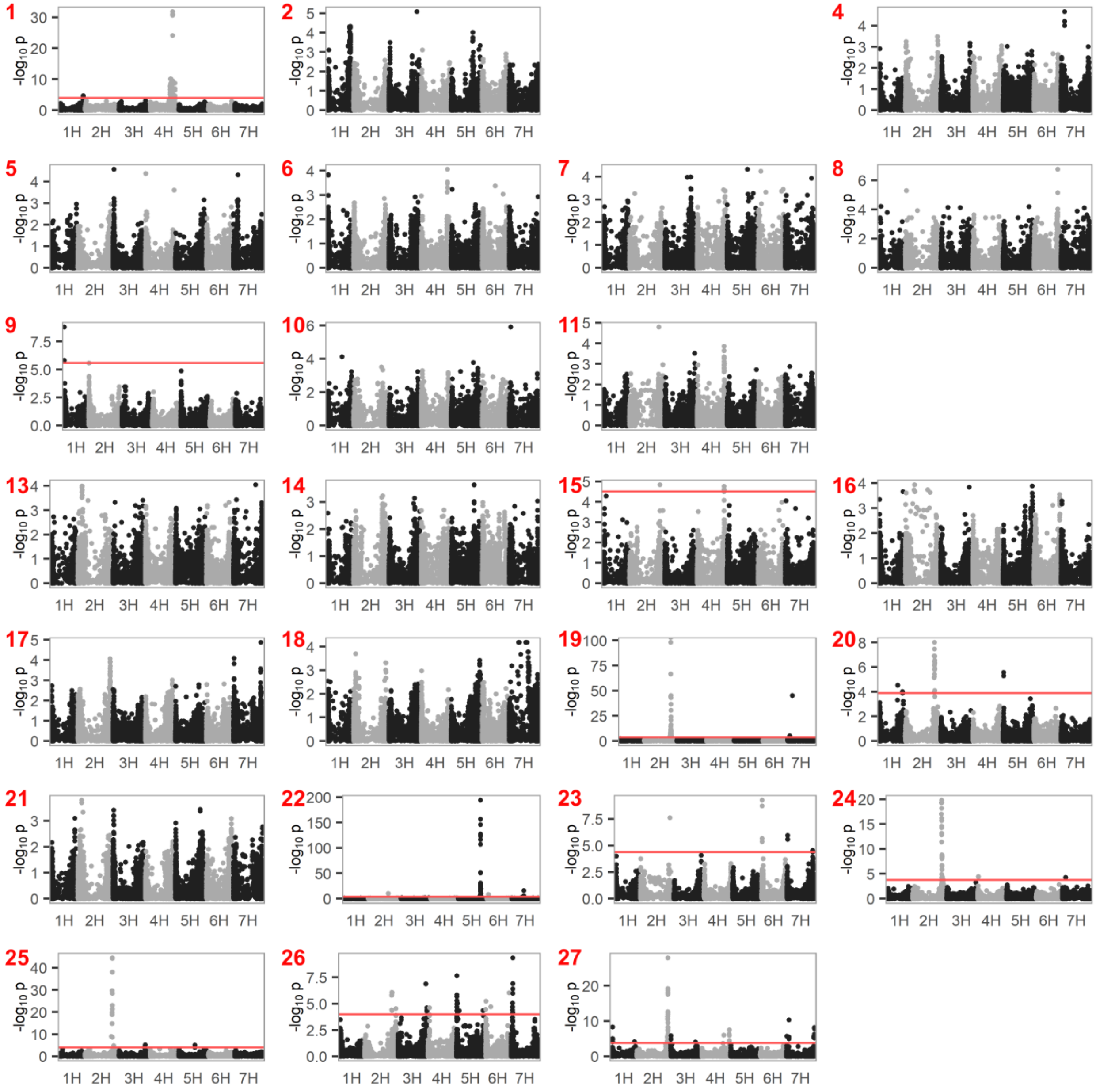

**Fig. S4. Manhattan plots of GWAS results from spring barley only dataset (n=370)**. Trait numbers are annotated at the top left corner of each Manhattan plot. Red horizontal line represents FDR = 0.05. If absent no significant marker was identified. Plots for trait 3, 12 and 28 are missing here because these traits are not segregating in the spring barley dataset.

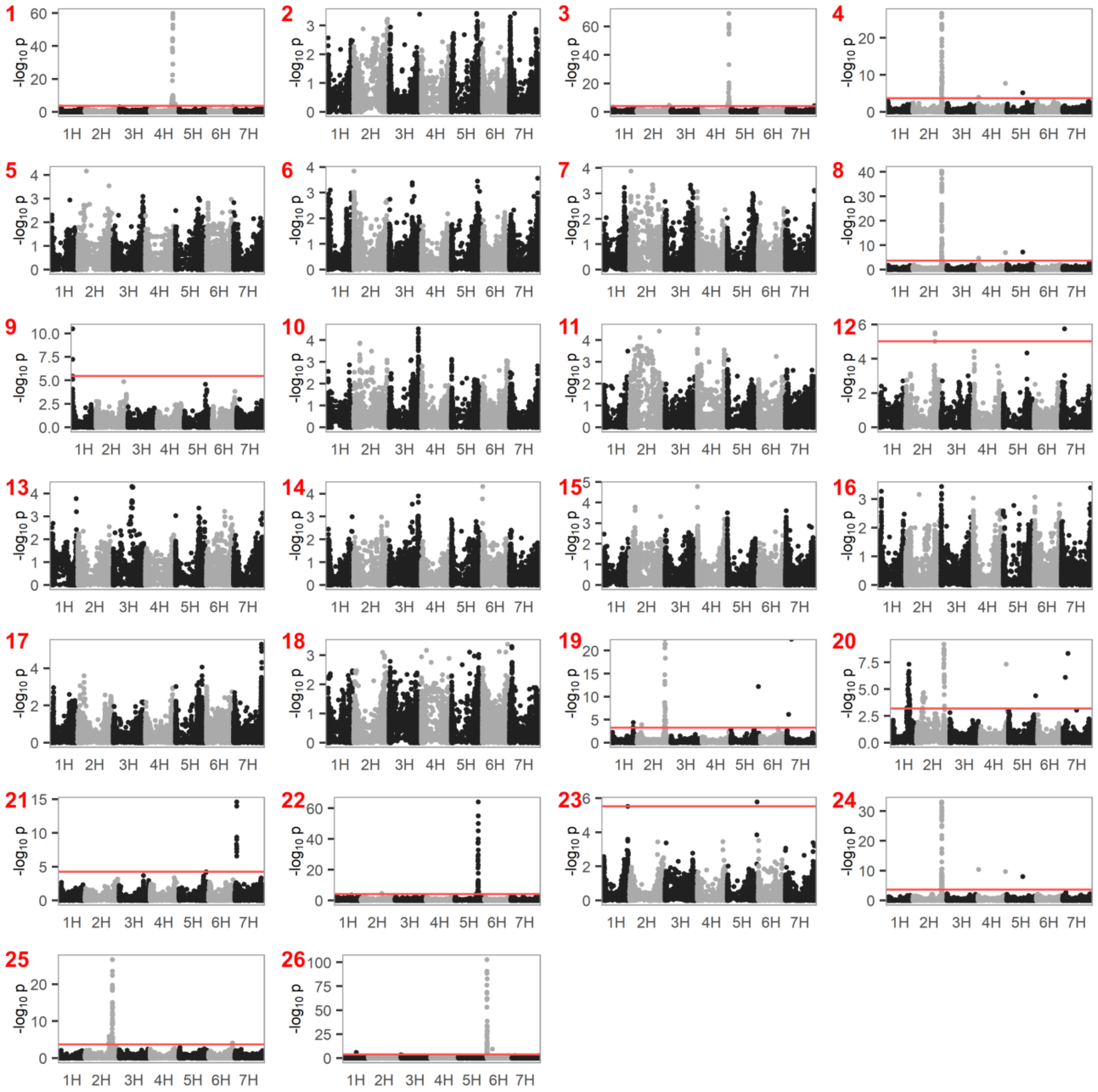


**Fig. S5. Manhattan plots of GWAS results from winter barley only dataset (n=335)**. Trait numbers are annotated at the top left corner of each Manhattan plot. Red horizontal line represents FDR = 0.05. If absent no significant marker was identified. Plots for trait 27 and 28 are missing here because these traits are not segregating in the winter barley dataset.
